# FGF9 treatment reduces off-target chondrocytes from iPSC-derived kidney organoids

**DOI:** 10.1101/2024.08.23.609318

**Authors:** Virginie Joris, Anika Schumacher, Paula Marks, Maria Eischen-Loges, Martijn van Griensven, Vanessa L.S. LaPointe

## Abstract

Renal failure due to drug nephrotoxicity or disease is frequently observed in patients. The development of *in vitro* models able to recapitulate kidney biology offers new possibilities to study drug toxicity or model diseases. Induced pluripotent stem cell–derived kidney organoids already show promise, but several drawbacks must be overcome to maintain them in culture, among which is the presence of non-renal cell populations such as cartilage. We modified the culture protocol and maintained kidney organoids in medium containing FGF9 for one additional week compared to the control protocol (Takasato). In comparison to the control, the FGF9-treated kidney organoids had reduced cartilage at day 7+25 and diminished chondrocyte marker expression. Importantly, the renal structures assessed by immunofluorescence were unaffected by the FGF9 treatment. This reduction of cartilage produces a higher quality kidney organoid that can be maintained longer in culture to improve their maturation for further *in vivo* work.

**Highlights:**

- Kidney organoids develop cartilage between days 7+18 and 7+25
- Extending the FGF9 supplementation reduces cartilage and chondrocyte markers
- FGF9-treated organoids present less EMT marker expression than control organoids
- Renal structures are not affected by the extended FGF9 treatment

**eTOC blurb:** LaPointe and colleagues showed that a one-week extension of FGF9 supplementation in iPSC-derived kidney organoids leads to a reduction of off-target cartilage. The renal structures are not impacted by FGF9 treatment and EMT markers are reduced. This modified organoid protocol will enable longer culture periods, a benefit for the use of organoids for screening or therapies.

## Introduction

Renal failure due to nephrotoxic drugs and diseases is frequently observed in patients and leads to a gradual loss of kidney function with a diminished glomerular filtration rate.^1^ The understanding of the mechanisms behind these processes is not completely elucidated and usually requires *in vivo* models. Nevertheless, *in vitro* models have the potential to revolutionize our ability to develop drugs and screen for potential toxic effects. Moreover, they also offer the possibility to develop disease models to study new therapeutic approaches.^2^ Indeed, a lot of effort has gone into making *in vitro* models that are able to recapitulate the biology of a functional organ. Using knowledge of developmental biology, human induced pluripotent stem cells (hiPSCs) can be directed to produce organoids mimicking various organ systems.^3^ Organoids present many advantages compared to 2D cultures, as they are 3D multicellular systems that are more physiologically representative and possess interactions between different cell types. Kidney organoids are a good example, as they can recapitulate renal structures as well as some of the cellular complexity of a human kidney and have demonstrated the potential to augment glomerular filtration upon transplantation *in vivo*.^4–6^ Their physiologic functionality *in vitro* demonstrates that kidney organoids might be suitable for studies usually requiring an *in vivo* model such as drug screening, drug discovery, or toxicity studies. This kidney organoid model may also open the door for the study of kidney development or diseases. Indeed, kidney diseases commonly involve the interactions of different cell types, and the multicellular aspect of kidney organoids allows their use as *in vitro* kidney disease models. As such, kidney organoids are already used to model some diseases, such as ciliopathies or polycystic kidney disease, and to study the mechanisms of renal pathologies as well as potential treatments.^7–9^

Many research efforts have been made to generate organoids that reproduce the key morphological and functional features of human kidneys. For example, while kidney organoids present renal structures observed in a native kidney, namely glomeruli, tubuli, and loops of Henle, they lack vascularization and phenotypic maturation. These drawbacks make the transposition of experiments on organoids to an adult kidney difficult. Several studies have addressed the problem of vascularization; for example, Low and colleagues employed a dynamic modulation of WNT signaling using CHIR to control the production of VEGFA and induce the development of a vascular network.^10^ In our lab, we demonstrated that hypoxic culture of kidney organoids induced upregulation of VEGFA-189 and -121 and downregulation of the antiangiogenic isoform VEGFA-165b, leading to an increase in vessel length.^11^ Furthermore, the use of decellularized kidney extracellular matrix during organoid transplantation led to increased vascularization by the host endothelial cells and improved maturation of the organoid after engraftment.^12^ Unfortunately, a persistent problem is that kidney organoids cannot be kept in culture for long periods without the undesired appearance of off-target cell populations, which worsens their maturation potential.

Off-target, non-renal cell populations represent 10–20% of the kidney organoid, appear around 18 days of culture, and mainly comprise neurons, myocytes and chondrocytes.^13^ Wu et al. first performed single-cell RNA sequencing and showed the expression of neural and muscular markers. Inhibition of these markers via a pharmacological approach reduced the neuron population development without affecting kidney structures^14^, establishing the possibility to improve kidney organoids using a small-molecule approach. The appearance of chondrocytes in kidney organoids is less studied as it requires maintaining the organoids in culture for a long time. Indeed, the majority of studies using kidney organoids focus on short timepoints and thus did not observe the appearance of this off-target population. Interestingly, the public transcriptomics data of Wu et al. also shows the appearance of a cluster of cells 19 days after aggregation expressing COL2A1 in kidney organoids cultured with the Takasato protocol. Moreover, the appearance of cartilage in kidney organoids was observed after *in vivo* engraftment of the organoid under the renal capsule of mice.^15,16^ The cartilage tissue appeared 4 weeks after graft transplantation and expanded over time. Transplantation of kidney organoids with decellularized kidney extracellular matrix resulted in a reduction of cartilage *in vivo* and emphasizes the importance of the microenvironment. In the study of Bantounas et al., the authors transplanted organoids composed of kidney progenitors in mice and still observed the appearance of cartilage weeks after the graft transplantation, highlighting the possible dedifferentiation of renal structures.^15^ The reason for the consistent observation of chondrocytes in kidney organoid culture and transplantation is not understood. Cartilage formation involves the condensation of mesenchyme tissue, which differentiates into chondrocytes and produces the extracellular matrix protein collagen 2 (COL2A1). The pathways leading to chondrocyte differentiation mainly involve the SOX protein family, particularly SOX9.^17^

To understand the link between chondrocytes and renal cells, we took inspiration from what happens in the adult kidney in the context of injury or stress where dedifferentiation of renal structures can occur. In the adult kidney, renal tissue can regenerate via the dedifferentiation of tubules, notably through the EGFR pathway.^18^ This pathway also induces a transitory increase of SOX9 expression, particularly in proximal tubules, and can last for 2–3 days to induce healing of the injured tissue.^19^ The transitory aspect of SOX9 expression is highly important, as this transcription factor is a key player in other processes including chondrogenesis.^20^ Indeed, during cartilage formation and maintenance, SOX9 secures the chondrocytic lineage, ensures cell survival and regulates genes implicated in cartilage structure.^21^

In the present study, we show that both SOX9 expression and protein levels increase during the same period and remain elevated until later time points, which correlates with the appearance of cartilage between days 18 and 25. With the aim of preventing the dedifferentiation of epithelial cells into mesenchymal cells during the development of the kidney organoid, we modulated the existing culture protocol. The growth factor FGF9, which is used in many protocols for the generation of kidney organoids, is important in normal kidney development^22,23^ but has also been shown to reduce the differentiation of human mesenchymal stem cells into cartilage.^24^ Therefore, we extended the incubation of kidney organoids with FGF9 from day 5 until day 12. We observed a clear reduction of the appearance of cartilage at the latter time point, which was correlated with a reduction of cartilage markers.

## Results

### Non-renal cell populations appeared between day 7+18 and 7+25 of kidney organoid culture

To induce the formation of kidney organoids from iPSCs, we used a protocol comprising 2D and 3D differentiation steps as illustrated in Figure 1A. We maintained the organoids in culture for up to 7+25 days. While organoids at day 7+18 presented a good overall shape (Fig. 1B), we observed that their shape became less regular by day 7+25 (Fig. 1C). Upon assessing the consequence of the prolonged culture on the development of renal structures, we observed that at day 7+18, the kidney organoids comprised glomeruli (NPHS1), tubules (ECAD), proximal tubules (LTL) (Fig. 1D) as well as loops of Henlé (SLC12A1) and the expected stromal population (MEIS) (Fig. 1E). By day 7+25, the kidney organoids still showed positive immunostaining for tubules, glomeruli, and loops of Henlé, but also showed the appearance of an off-target population absent of renal markers (asterisks in Fig. 1F and 1G).

**Figure 1:**
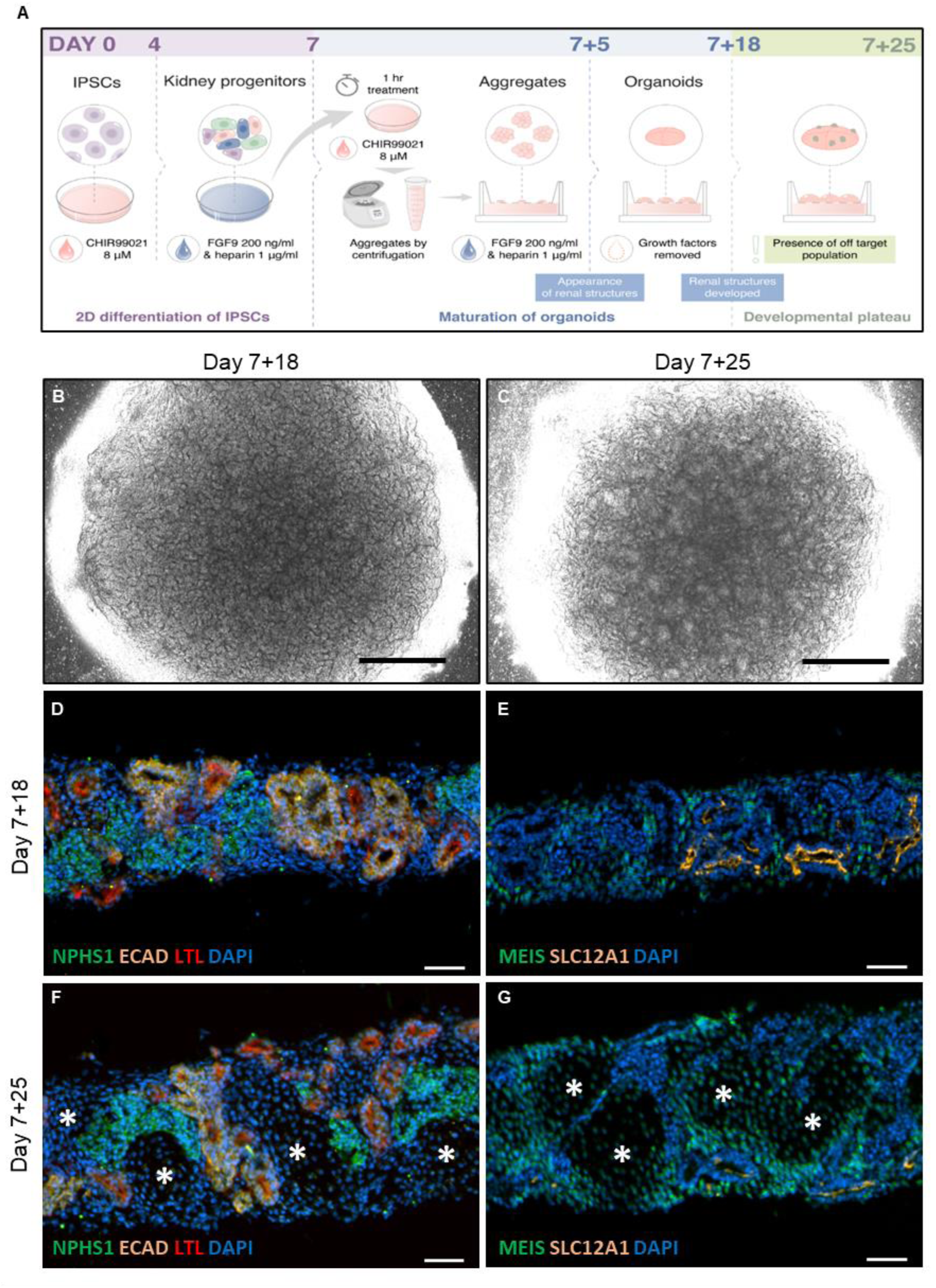
Off-target cell population develops in iPSC-derived kidney organoids and disrupts the renal structures’ development. (A) Schematic of kidney organoid culture. iPSCs were stimulated with CHIR99021 for 3 days and a FGF9/heparin cocktail for 4 days. After a 1 h pulse of CHIR99021, cells were aggregated and cultured at the air–liquid interface for 5 days in the presence of the FGF9/heparin cocktail. Organoids were then cultured until day 7+25 without growth factors. A spherical kidney organoid shape was observed at day 7+18 (B) and 7+25 (C) by brightfield microscopy. (D–G) Renal structures assessed by immunofluorescence in cryosections show (D, F) glomeruli (NPHS1), proximal tubules (LTL), distal tubules (ECAD); (E, G) loops of Henlé (SLC12A1) and a stromal population (MEIS1/2/3). Nuclei are stained with DAPI. Asterisks indicate an off-target cell population at day 7+25 (F, G). Scale bars represent 1000 µm (B–C) and 50 µm (D–G).

### The non-renal cell population was identified as cartilage progressively developing over time in culture

As described previously in the literature^15,16^, we showed that the predominant off-target cell population observed at day 7+25 is cartilage. While Alcian blue staining of the whole kidney organoid and cryosections did not show cartilage in organoids at day 7+18 (Fig. 2A and 2B), by day 7+25 the staining revealed abundant cartilage in the non-renal cell population (Fig. 2C and 2D). This observation was correlated with increased expression of the cartilage-related transcripts, *SOX9* (*p* < 0.05), *COL2A1* (*p* < 0,001), *ACAN* (*p* < 0.01), *COL1A*1 (*p* < 0.001), and *COL10A1* (*p* < 0.01) in day 7+25 kidney organoids measured by qPCR (Fig. 2E–I). Some of these markers, such as COL2A1 and SOX-9, were also assessed at the protein level using western blotting (Fig. 2J). COL2A1 protein level presented a 3.5 times increase (*p* < 0.01) at day 7+25 compared to day 7+18 (Fig. 2J and 2L), whereas SOX-9 levels remained unchanged (Fig. 2J and 2K).

**Figure 2:**
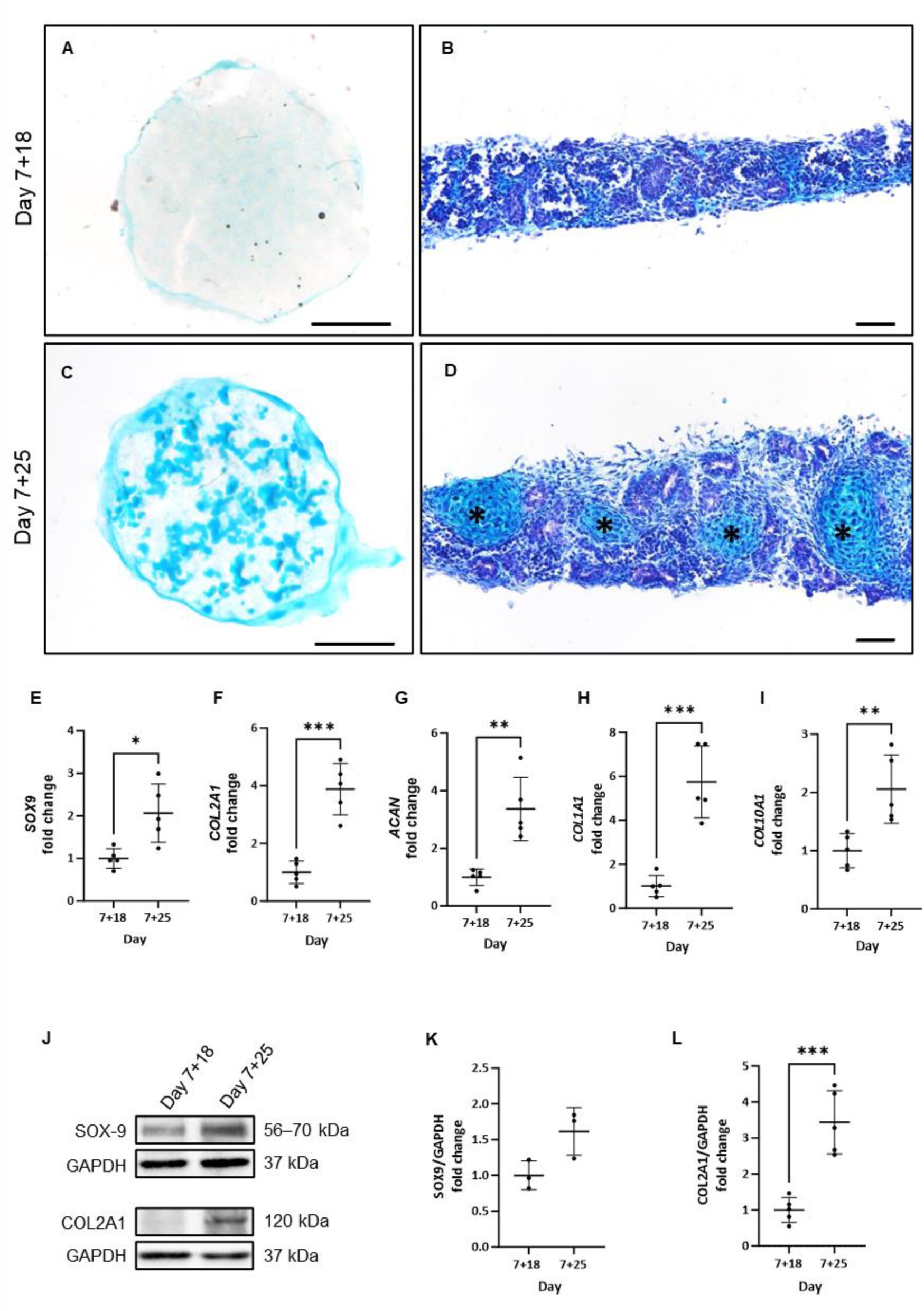
The off-target population was identified as cartilage developing between days 7+18 and 7+25. (A–D) Alcian blue staining revealed the presence of cartilage at days 7+18 and 7+25 in the whole organoid (left images, scale bars represent 1000 µm) and cryosections (right images; scale bars represent 50 µm). (E–I, N=5) Progressively and significantly increased expression of five different markers of chondrogenesis was detected by qPCR at day 7+18 and 7+25. \**p* < 0.05; \*\**p* < 0,01; \*\*\**p* < 0.001 based on -fold change relative to day 7+18. (J) Western blotting of SOX9 and COL2A1 protein levels in kidney organoids from day 7+18 to 7+25. GAPDH levels are shown as loading controls. (K–L) Quantification of protein levels using ImageJ normalized to GAPDH showed a significant increase (\*\*\**p* < 0.001; N=5) of COL2A1 (L) from day 7+18 to 7+25. SOX9 levels (K; N=3) did not show significant differences.

### At day 7+25, cartilage did not form in FGF9-treated kidney organoids

With the aim to prevent the cartilage formation detected at day 7+25, FGF9 without heparin was added from day 7+5 until day 7+12 (Fig. 3A). Organoids that underwent this treatment are referred to as FGF9-treated organoids (day 7+25+FGF9), while the organoids cultured in regular conditions are referred to as control organoids (day 7+25). Using Alcian blue to stain the whole organoids and cryosections, we observed that the cartilage present in control organoids (Fig. 3B and 3C) was absent in the FGF9-treated organoids at day 7+25 (Fig. 3D and 3E). At the molecular level, *SOX9*, *ACAN*, *COL2A1*, *COL1A1*, and *COL10* mRNA expression were also decreased (Fig. 4A–E, *p* < 0.01). A three times decrease of SOX9 protein levels was also observed in FGF9-treated organoids compared to control organoids at day 7+25 (Fig. 4F and G, *p* < 0.01), associated with a four times decrease of COL2A1 protein levels (Fig. 4F and H, *p* < 0.01).

**Figure 3:**
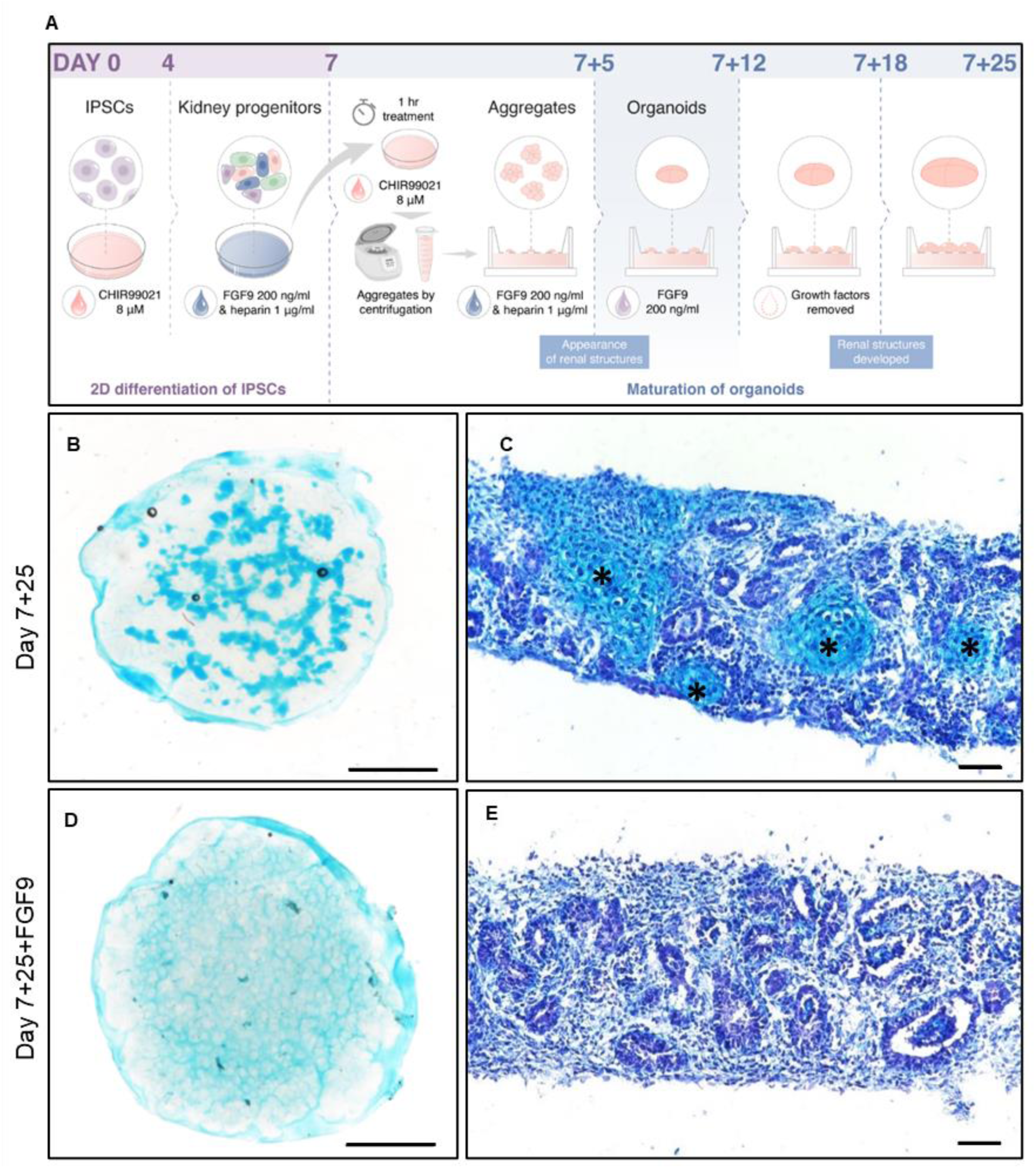
FGF9 treatment abrogates cartilage formation at day 7+25. (A) Schematic of the FGF9 treatment, which was extended after aggregation from day 7+5 to day 7+12. (B–E) Cartilage stained with Alcian blue in whole organoids (left images, scale bars represent 1000 µm) and on cryosections (right images, scale bars represent 50 µm) was strongly reduced with FGF9 treatment (bottom row).

**Figure 4:**
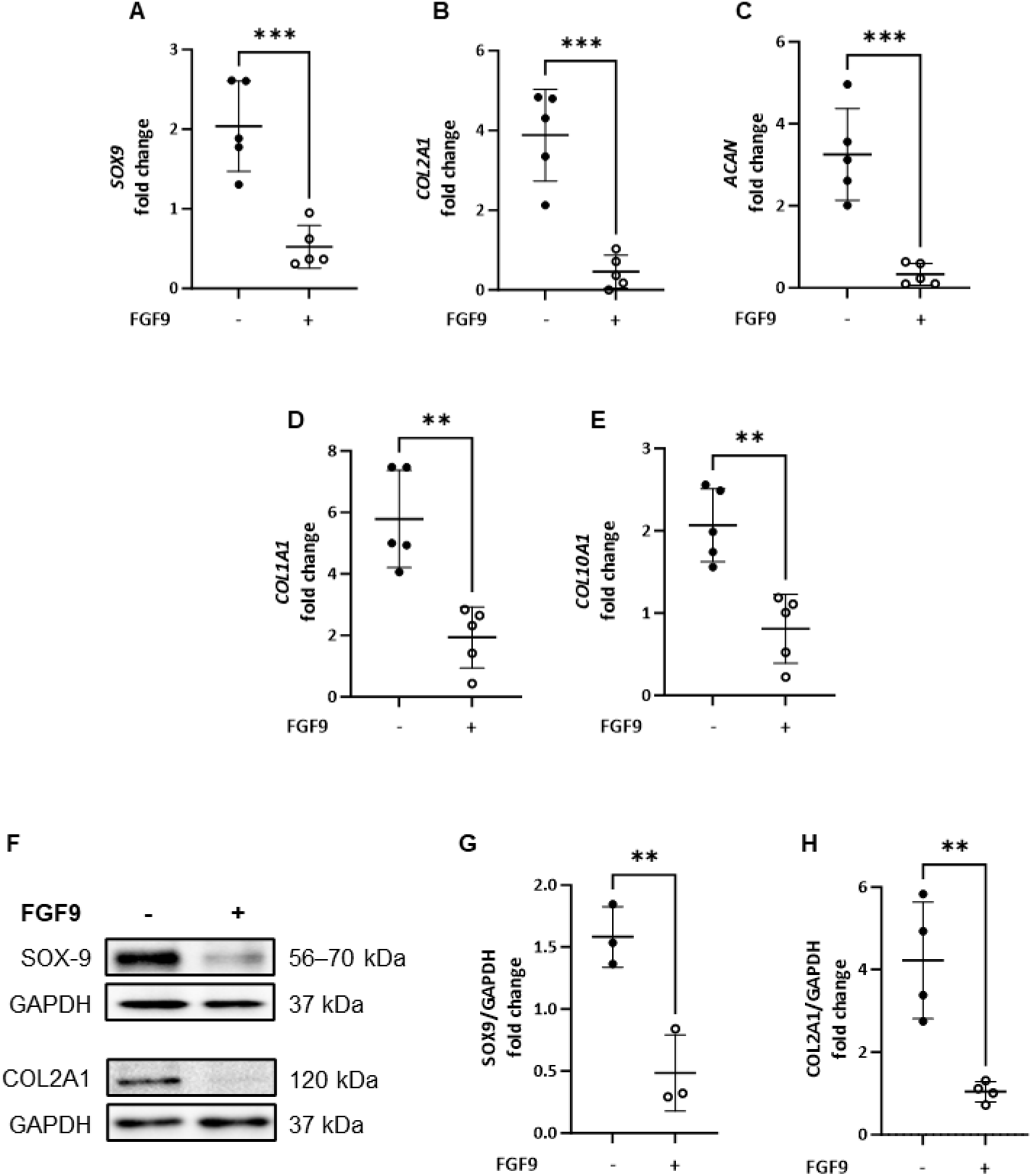
FGF9 treatment reduces expression of cartilage markers at day 7+25. (A–E) FGF9 treatment significantly decreased five markers of chondrogenesis in kidney organoids at day 7+25 compared to control organoids. Gene expression was assessed by qPCR and shown as -fold change compared to expression at day 7+18. \*\**p* < 0.01; \*\*\**p* < 0.001; N=5. (F) Western blotting of SOX9 and COL2A1 protein levels in kidney organoids at day 7+25 showed decreased expression in FGF9-treated organoids compared to untreated controls. GADPH levels are shown as loading controls. (G–H) Quantification of protein levels confirmed significantly decreased expression of SOX9 (G) and COL2A1 (H) at day 7+25 with FGF9 treatment (+) compared to nontreated (–) organoids. \*\**p* < 0.01 from 3–4 samples. Relative expression was assessed as the -fold change compared to day 7+18.

### FGF9-treated organoids correctly developed renal structures

To ensure the FGF9 treatment had no negative impact on the development of renal structures in kidney organoids, cryosections were stained for renal markers (Fig. 5A–D): NPHS1 (glomeruli), ECAD (distal tubules), LTL (proximal tubules) and SLC12A1 (loops of Henlé). Unlike the control organoids, we found that FGF9-treated organoids at day 7+25 possessed typical renal structures (Fig. 5C and 5D). Indeed, specifically SLC12A1 was hardly present by day 7+25 but the FGF9 treatment restored it to levels similar to day 7+18. This observation correlated with the expression of the renal structures’ markers measured by qPCR, showing no detrimental impact of the FGF9 treatment in organoids at day 7+25 (Fig. S1). This result suggests that the phenotype of FGF9-treated organoids at day 7+25 is close to the one observed in control organoids at day 7+18.

**Figure 5:**
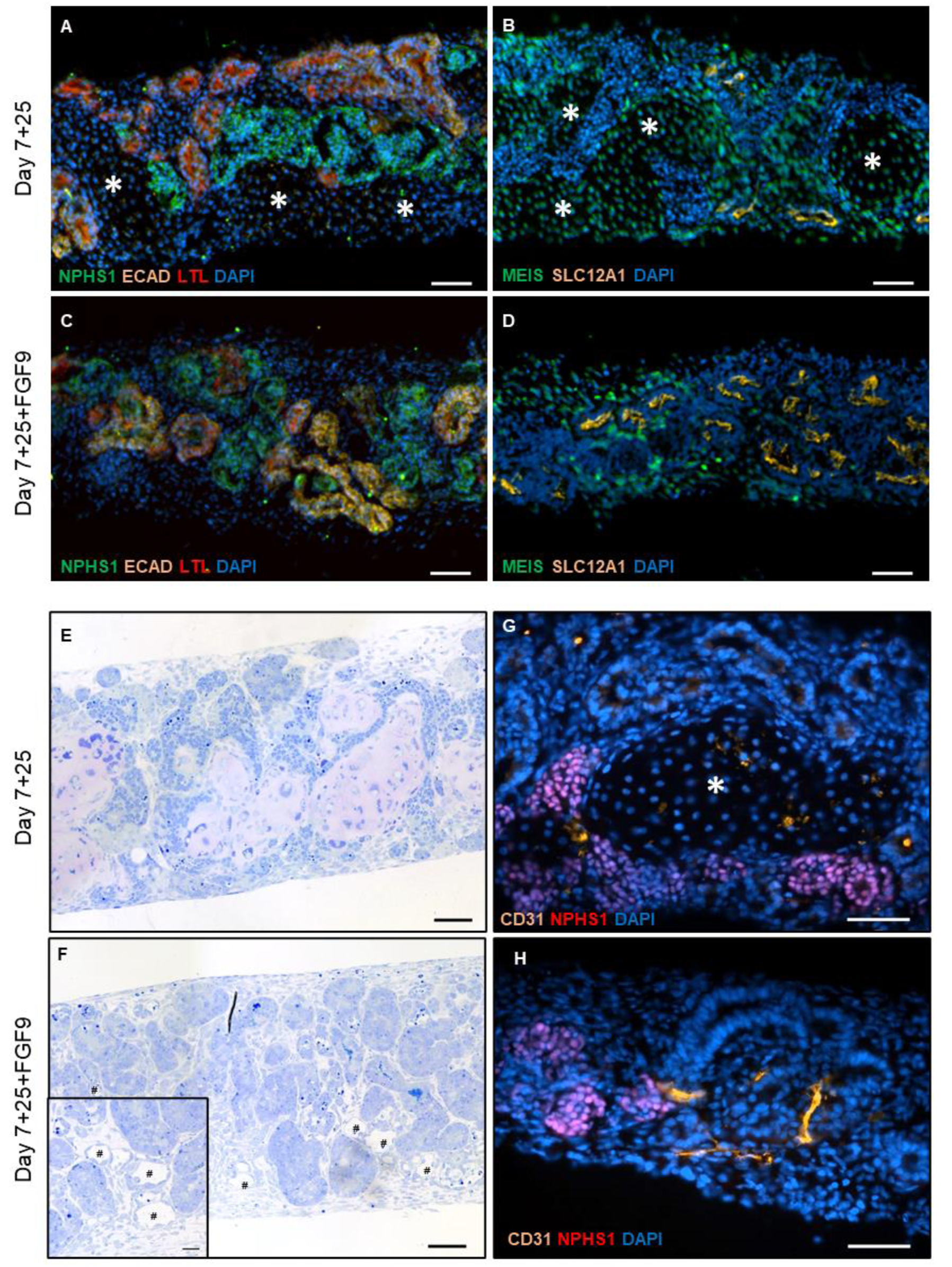
FGF9-treated organoids show renal and vessel-like structures. (A–D) Renal structures assessed by immunofluorescence in cryosections show (A, C) glomeruli (NPHS1), proximal tubules (LTL), distal tubules (ECAD); (B, D) loops of Henlé (SLC12A1) and stromal population (MEIS1/2/3). Nuclei are stained with DAPI. The off-target cell population (asterisks) observed in day 7+25 untreated organoids (top row) was not observed with FGF9 treatment (bottom row). Scale bars represent 50 μm. (E–F) Toluidine blue staining showed the appearance of vessel-like structures (#) in FGF9-treated organoids (F) that were absent in control organoids (E) at day 7+25. Scale bars represent 50 µm; inset represents 20 µm. (G–H) Immunostaining of cryosections showed an increase of the endothelial marker CD31 in FGF9-treated organoids (H) compared to controls (G). Scale bars represent 50 µm.

### FGF9-treated kidney organoids developed vessel-like structures

While control organoids do not contain vessels at day 7+25 (Fig. 5E), FGF9-treated organoids developed vessel-like structures, as observed by toluidine blue staining (Fig. 5F). This observation was correlated with an increased level of the vascularization marker CD31 (PECAM1) in immunofluorescence (Fig. 5G and 5H) and *PECAM1* expression measured by qPCR (Fig. S2) in FGF9-treated organoids compared to control organoids.

### FGF9-treated kidney organoids presented a complex phenotype and albumin uptake

We assessed the level and location of the water channel aquaporin 2 (AQP2) in control (Fig. 6A–B) and FGF9-treated organoids (Fig 6C–D). We observed that the AQP2 staining was higher with FGF9 compared to the control and located in tubules. Moreover, control (Fig. 6E–F) and FGF9-treated organoids (Fig. 6G–H) were incubated for two days with Alexa Fluor 647-bovine serum albumin (BSA). We observed more BSA located in the tubules of the FGF9-treated organoids compared to the control.

**Figure 6:**
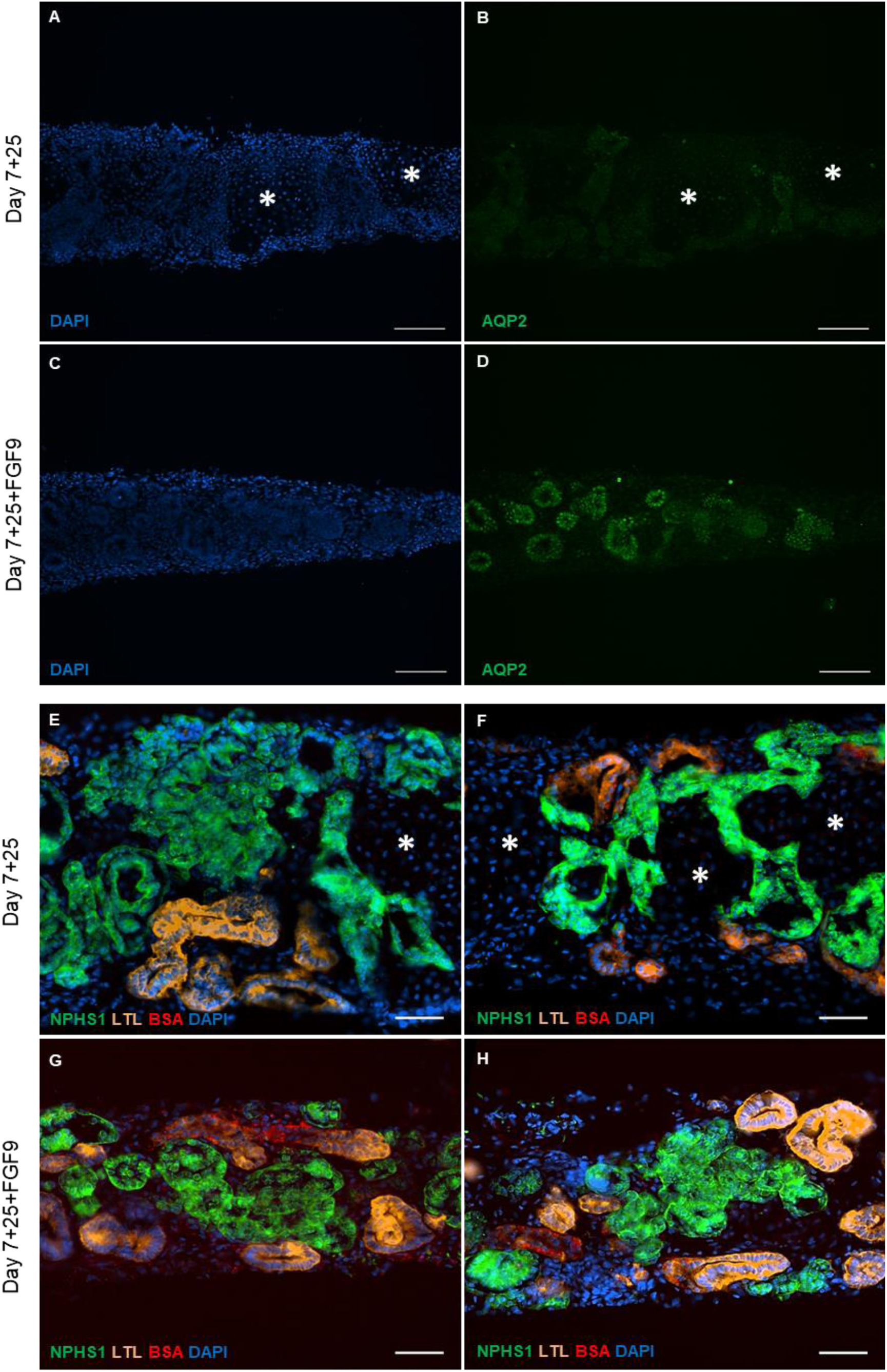
FGF9-treated organoids present a more complex phenotype. (A–D) Immunostaining of cryosections showed an increase of the vasopressin-induced water channel aquaporin 2 (AQP2) in FGF9-treated organoids. Nuclei are stained with DAPI. Scale bars represent 100 µm. The off-target cell population (asterisks) observed in day 7+25 untreated organoids was not observed with FGF9 treatment. (E-H) BSA uptake assay revealed a higher staining (red) in FGF9-treated organoids compared to control at day 7+25. Nephrons (NPHS1) and proximal tubules (LTL) are present in both conditions. Nuclei were stained with DAPI. Scale bars represent 50 µm.

### FGF9-treated organoids at day 7+25 showed low levels of epithelial to mesenchymal transition (EMT)

We assessed the levels of EMT markers vimentin and α-SMA at days 7+5, +10, +14, +18, +25 and +25+FGF9. We observed that EMT markers increased from day 7+5 until day 7+25 in control organoids. FGF9-treated organoids at day 7+25 had levels of vimentin and α-SMA similar to day 7+5, showing that EMT was halted with the treatment (Fig. S2A–C).

### Cartilage was found in FGF9-treated organoids at day 7+32

To assess if the FGF9 treatment allows kidney organoids to be kept longer in culture with no off-target appearance, we maintained control and FGF9 organoids in culture for 32 days after aggregation (7+32). In the control organoids, we observed that cartilage formation continued to increase from day 7+25 to day 7+32 (Fig. S3A and S3B). In the FGF9-treated organoids at day 7+32, we did not observe the appearance of large islands of cartilage in Alcian blue–stained whole organoids (Fig. S3C), but we did notice small islands of cartilage at the organoid edges (Fig. S3D), albeit less than control organoids at day 7+25. Moreover, the expression of *SOX9* (*p* < 0.001), *COL2A1* (*p* < 0.001), *ACAN* (*p* < 0.001), and *COL10A1* (*p* < 0.001) (Fig. S3E–I) was decreased in FGF9-treated organoids compared to control organoids. These findings were confirmed by COL2A1 protein levels in western blots (Fig. S3J).

## Discussion

The possibility of creating kidney organoids derived from human iPSCs already improves the understanding of kidney development and response to drugs. Moreover, organoids also seem to be a promising therapeutic approach as they could be used for transplantation into patients suffering from renal failure. However, despite their ability to recapitulate renal structures and function, several drawbacks prevent their long-term use for developmental and toxicity studies, as well as their use as a therapy. For example, the emergence of off-target differentiation products such as cartilage remains a major issue as it profoundly disrupts organoid structure. In this study, we aimed to adjust the differentiation protocol developed by Takasato e*t al*. in order to prevent the appearance of cartilage *in vitro* ^4^. We found that extending the FGF9 incubation from day 7+0 to day 7+12 after aggregation at the air–liquid interface significantly reduced the off-target cartilage until day 7+25. This presence of cartilage was assessed with Alcian blue staining and the measurement of cartilage markers, such as SOX9 and COL2A1, in qPCR and western blot. Importantly, this modified protocol did not negatively affect renal structures as evaluated by immunofluorescence and PCR. Indeed, the organoids still presented nephron and tubular structures, with an increased number of vessel-like structures.

We observed that the appearance of cartilage in kidney organoids occurred between day 7+18 and day 7+25 and was correlated to an increased expression of *ACAN*, *COL2A1*, and *SOX9*. These results correlate with the appearance of a cluster of cells expressing *COL2A1* in the previous database of single-cell transcriptomics reported by Wu *et al*.. Additionally, Nam *et al*. previously reported that human iPSC-derived kidney organoids grafted under the murine renal capsule developed cartilage islands 4 weeks post-transplantation. Interestingly, the cartilage islands enlarged 6 weeks after transplantation.^15^ In our *in vitro* organoids, we observed a similar development albeit at a different time scale; at day 7+32, the cartilage invaded a larger area than at day 7+25, was associated with a higher expression of cartilage markers, and slowly took over the other cell populations over time. In their study, the authors also showed that the use of a decellularized kidney extracellular matrix did not remove, but reduced the appearance of cartilage, demonstrating the importance of the environment in the development of kidney organoids. They suggested that non-kidney precursor cells were present in the organoids and developed into cartilage over time. However, the transplantation of pre-selected kidney progenitor cells under the renal capsule still led to the formation of cartilage, instead suggesting a dedifferentiation of renal structures and their redifferentiation into cartilage.^16^ We hypothesize that dedifferentiation may be occurring in our organoid culture.

Indeed, dedifferentiation is already described in the adult kidney. Upon injury or stress, EGFR activation leads to the dedifferentiation of renal cells through the activation of the MAPK/ERK and PI3K pathways.^18^ This activation also induces a transitory increase of SOX9 expression, particularly in proximal tubules, and can last for 2–3 days to induce regeneration of the injured tissue.^19^ Interestingly, in our previous work ^25^, we showed that the EGFR pathway is activated around day 7+10 in the control organoids. Moreover, preliminary results showed an increase of SOX9, a key player in cartilage formation, early in the formation of kidney organoids, around day 7+10 and its level was maintained until day 7+ 25 (data not shown). Of course, the kidney organoids produced *in vitro* do not possess the same properties as the adult kidney, as they are developmentally comparable to first-trimester embryonic kidneys ^26^. This immature environment may not allow SOX9 expression to be controlled correctly, leading to a sustained increase of this transcription factor and the subsequent increase of COL2A1 expression and cartilage formation. Therefore, whether controlling SOX9 expression also ameliorates cartilage formation in kidney organoid cultures could be explored in subsequent studies.

To prevent the probable dedifferentiation of the tubules, we extended the FGF9 treatment from day 7+0 until day 7+12 instead of day 7+5, as in the original protocol. Along with FGF20, FGF9 is an important growth factor for kidney development and stemness of renal progenitor cells.^22^ Interestingly, expression of FGF9 in the kidney is localized to ureteric buds and nephron progenitor cells. Its deletion leads to a drastic reduction of nephron progenitor cells and, *in fine*, to kidney agenesis.^27^ On the other hand, FGF9 is described as a positive trigger in cartilage development. Therefore, it was counterintuitive to use it in our kidney organoids to remove cartilage. However, FGF9 can also play a developmental stage–dependent role in cartilage formation. For example, in stem cells with chondrogenic potential, the addition of FGF9 in the early phase of chondrogenic differentiation prevented cartilage formation.^24,28^ The expression of receptors FGFR1 and FGFR3 are important for this response to FGF9. At an early stage, stem cells treated with FGF9 showed unchanged FGFR1 expression, but reduced FGFR3 expression. In contrast, in the late phase of differentiation, both receptors’ expression was increased after FGF9 treatment, leading to more cartilage deposition. Therefore, in these cells, the effect of FGF9 on ECM deposition is mediated by FGFR3 and the FGFR3/FGFR1 ratio and is a preset to cartilage appearance.^24^ Conversely, a potential downregulation of FGFR3 due to FGF9 treatment in renal cells was not expected to have an impact. Indeed, in the kidney, a downregulation of FGFR3 was shown to produce no adverse effects on renal development.^23^ However, the overactivation of FGFR3 is known to induce EGFR activation and EMT.^29^

EMT is already known in kidney organoids and is associated with partial dedifferentiation and increased expression of stemness markers.^5,30^ Previous work in our lab showed that EMT markers such as vimentin, α-smooth muscle actin (αSMA), collagen 1a1, and collagen 6a1 are increased during kidney organoid development using the Takasato protocol.^5,31^ We observed that FGF9 treatment reduced levels of vimentin and α-SMA (Fig. 6), improving the global quality of the organoids at day 7+25.

In addition to the reduction of cartilage at day 7+25, we noticed that FGF9-treated kidney organoids also present vessel-like structures that were not found in control organoids. FGF9 is known to enhance angiogenesis during long bone repair. FGF9^-/+^ mice present less VEGFA and PECAM1 than wild-type mice and showed impaired angiogenesis during bone healing.^32^ Moreover, the addition of FGF9 during angiogenesis enhanced the production of vasoresponsive microvessels.^33^ In this work, we suspect that endothelial cells formed during the kidney differentiation, and that FGF9 increased their appearance. The presence of these vessel-like structures can help with future ambitions regarding engraftment *in vivo*. Specifically, the presence of a vascular network in the organoid could help its perfusion by the host vessels.

Lastly, we also observed a higher expression of aquaporin 2 in tubules of FGF9-treated organoids compared to control as well as improved uptake of BSA. Aquaporin 2 is mainly located in the connecting and collecting tubules in the adult kidney, where it is responsible for vasopressin-induced water reabsorption.^34^ Interestingly, a dysregulation of aquaporin 2 is observed in renal failure or diabetic nephropathy. ^34,35^ More recently, aquaporin 2 expression was also studied in proximal tubules where its overexpression reduced the cell damage due to ischemia-reperfusion. ^36^ The mechanism by which FGF9 modulates aquaporin 2 level is not clear but might involve activation of PI3K or PKA. ^34,37^ This upregulation of aquaporin 2 and BSA uptake reflect a more complex renal phenotype in FGF9-treated organoids than in control at day 7+25.

Longer culture of FGF9-treated organoids until day 7+32 showed small islands of cartilage reappearing (supp figures), albeit in much smaller quantities than in the control organoids at day 7+25. FGF9 is known to maintain the stemness of the nephron progenitors and might therefore just delay the appearance of cartilage due to a delay in the differentiation–dedifferentiation process.^22^ While this type of improvement is long enough for some purposes such as disease modeling or drug testing, it is possibly not long enough for regenerative medicine.

To conclude, we optimized the culture protocol of kidney organoids by extending the time of incubation with FGF9 after aggregation at the air–liquid interphase. We managed to reduce the appearance of cartilage in organoids after 7+25 days of culture without affecting the renal structures. We also observed the appearance of more vessel-like structures. Since the cartilage began to reappear on day 7+32, we are aware that these organoids still need to be improved, maybe with the combination of other small molecules to maintain a renal phenotype over time and move them one step closer to the clinic. Moreover, the engraftment of these FGF9-treated organoids must also be performed to see if a renal phenotype can be maintained in an *in vivo* environment. For the moment, these FGF9-treated organoids, with reduced off-target cell populations and corresponding side effects, are of higher quality and can be used for disease modeling and drug testing.

## Material and methods

### Resource availability

Upon publication, all data associated with this manuscript will be made publicly available or upon request from the corresponding author (V. LaPointe).

### Culture and maintenance of human induced pluripotent stem cells (hiPSCs)

The hiPSC line LUMC0072iCTRL01 was provided by the hiPSC core facility at the University Medical Center Leiden (the Netherlands). Cells are cultured in 6-well plates coated with vitronectin (Fisher Scientific) using E8 medium (Thermo Fisher Scientific) supplemented with 1% penicillin–streptomycin (Gibco). The medium was refreshed daily and the cells were passaged twice weekly using TrypLE dissociation reagent (Invitrogen). After passaging, cells were seeded in the presence of RevitaCell (Fisher Scientific) overnight. Pluripotency and karyotype of the iPSCs were verified as described previously ^5^. All the key findings were repeated in two others hiPSC lines LUMC0099 and LUMC0031 provided by the UMC Leiden.

### iPSC-derived kidney organoid generation and treatment

Kidney organoids were generated at the air–liquid interface using a previously described protocol ^11^. Briefly, 10^5^ cells were seeded in a vitronectin-coated 6-well plate. After 24 h, regular E8 medium was replaced by STEMdiff APEL2 medium (STEMCELL Technologies) supplemented with 1% protein-free hybridoma medium II (PFHMII) (Fisher Scientific), 1% antibiotic–antimycotic (AA), and 8 µM CHIR99021 for 4 days to induce differentiation. This step was followed by an incubation with 200 ng/mL fibroblast growth factor 9 (FGF9, R&D Systems), and 1 µg/ml heparin (Sigma-Aldrich) for 3 days. After these 7 days of differentiation in 2D (denoted as day 7+0), cells were treated with CHIR99021 for 1 h, trypsinised and counted. 5×10^5^ cells were centrifuged at 425 RCF to form aggregates and placed on polyester membrane cell culture inserts with 1 µm pores (CellQART) to create a 3D culture at the air–liquid interface. The organoids were cultured in complete STEMdiff APEL2 medium supplemented with 200 ng/ml FGF9 and 1 µg/ml heparin for 5 additional days (denoted as day 7+5), after which the cocktail of growth factors was removed (Fig. 1A). Kidney organoids were then cultured until day 7+25 or day 7+32 in STEMdiff APEL2 medium supplemented with 1% PHFMII and 1% AA at 37 °C and 5% CO_2_.

To assess the impact of a prolonged FGF9 treatment on the organoids, 200 ng/ml FGF9 (without the simultaneous addition of heparin) was added from day 7+5 to day 7+12. Beginning on day 7+12, the organoids were maintained in complete STEMdiff APEL2 medium supplemented with 1% PHFMII and 1% AA until day 7+25 or day 7+32.

### Alcian blue staining of whole kidney organoids

To assess the onset of cartilage, the kidney organoids were fixed on days 7+18, 7+25 or 7+32 on the cell culture inserts in 70% ethanol overnight (ON) at 4°C. They were then incubated at room temperature (RT) for 1 h in 95% ethanol followed by acetone overnight. Alcian blue (0.03% (w/v)) diluted in 20% acetic acid was then added to the organoids for 4 h at RT followed by a rinse in 1% potassium hydroxide (KOH) (w/v) until the tissue was transparent. The organoids were stored in glycerol at 4°C before imaging on a Nikon SMZ25 automated stereomicroscope equipped with a 1× objective and a PHOTONIC LED-Set Ringlight stereomicroscope (SMZ25, Nikon). Z stack images were taken every 15 µm, images were analyzed using NIS-Elements software (version 5.30.06, Nikon).

### Gelatin/sucrose embedding of organoids for cryosections

On days 7+18, 7+25, or 7+32, kidney organoids were incubated for 20 min in 2% (v/v) paraformaldehyde diluted in phosphate-buffered saline (PBS) at 4°C. After a rinse in PBS, organoids were incubated in 15% (w/v) sucrose solution (in 0.1 M phosphate buffer pH 7.4) at 4°C for 24 h under agitation. The organoids are then transferred for 48 h into a 30% (w/v) sucrose solution at 4°C and set in a cryomold with the freezing buffer (7.5% (w/v) gelatin and 15% (w/v) sucrose in 0.1 M phosphate buffer). After letting the freezing medium harden on ice, the organoids were snap-frozen in liquid nitrogen. Cryosections were cut 16 µm thick on a CM3050S cryostat (Leica), mounted on Superfrost slides (Fisher Scientific), and used for immunostaining or Alcian blue staining. To remove the gelatin/sucrose embedding before any staining, cryosections were placed in PBS for 15 min at 37°C.

### Alcian blue and Toluidine Blue staining on sections

After removal of embedding and a quick wash in distilled water, the sections were incubated for 3 min in 3% (v/v) acetic acid diluted in water followed by incubation in a 1% (w/v) Alcian blue diluted in 3% acetic acid for 30 min at RT protected from light. After washing, samples were stained for 3 min with a nuclear fast red solution (Sigma-Aldrich). Slides were dehydrated and mounted in Ultrakitt mounting medium (VWR). For Toluidine Blue, ultrathin sections are dried and incubated for 30 s with one drop of staining consisting of 1% Toluidine Blue and 1% Sodium tetraborate diluted in milli-Q water. After a rinse, slides are mounted. Imaging was performed on an inverted Nikon Ti-S/L100 microscope, equipped with a Nikon DS-Ri2 camera using a CFI Plan Apochromat K 20×objective (NA: 0.75, WD: 1.0). Images were analyzed using NIS-Elements software (version 5.30.06, Nikon).

### Immunofluorescence

After embedding removal and in order to assess the renal structures, the slides were incubated for 20 min at RT in a blocking buffer containing 10% bovine serum albumin (BSA), 0.1 M glycine and 0.2% Tween-20 in PBS. Primary antibodies (Supplementary Table 1), diluted in PBS with 1% BSA, 0.1 M glycine and 0.2% Tween-20, were incubated overnight at 4°C. After rinsing in PBS with 0.2% Tween-20, the slides were incubated with secondary antibodies diluted in PBS with 1% BSA, 0.1 M glycine and 0.2% Tween-20 for 1 h at RT. Nuclei were stained using 4′,6-diamidino-2-phenylindole (DAPI, 0.1 μg/ml) in PBS with 0.2% Tween-20 for 5 min. Slides were mounted using Dako fluorescence mounting medium (Agilent Technologies).

BSA uptake staining was performed by incubating kidney organoids with an Alexa fluor 647–labelled BSA for two days at a concentration of 10 µg/ml. The organoids were then fixed, embedded, and cut as previously described. Slides were stained with LTL, mounted, and imaged.

For all the staining, imaging was performed on an automated inverted Nikon Ti-E microscope, equipped with a Lumencor Spectra light source, an Ador Zyla 5.5 sCMOS camera, and an MCL NANO Z200-N TI z-stage. The objective used was CFI PLAN APO LBDA 10× 0.45/4 mm. Images were analyzed using NIS-Elements software (version 5.30.06, Nikon).

### Western blot

Proteins were extracted from the kidney organoids using TRIzol (Invitrogen) as described by the manufacturer. After performing a bicinchoninic acid protein assay (Pierce), 15 µg of protein was loaded per well of an 8% or 10% acrylamide gel. Migration was performed at 120 V in a migration buffer (pH 8.3) consisting of 25 mM Tris-Base, 192 mM glycine, and 0.1% sodium dodecylsulfate (SDS) (Bio-Rad). Transfer to nitrocellulose membrane was performed at 4°C for 90 min in a transfer buffer consisting of 50 mM Tris, 40 mM glycine, and 1.5 mM SDS at a constant amperage of 350 mA. The membranes were then incubated in a blocking solution consisting of Tris-buffered saline (TBS) added with 5% BSA and 0.1% Tween-20 for 1 h at RT. Primary antibodies diluted in blocking solution were incubated overnight at 4°C. Primary antibodies used were the following: anti-SOX-9 (1/1000, Cell Signaling Technology, 82630), anti-GAPDH (1/10000, Cell Signaling Technology, 2118), anti-vimentin (1/500, Thermo Fisher Scientific, MA5-16409), anti-α-SMA (1/1000, Cell Signaling, 19245S) and anti-COL2A1 (1/1000; Abcam, ab34712). Membranes were rinsed twice in TBS with 0.1% Tween and incubated for 1 h with secondary antibodies coupled to a horseradish peroxidase (HRP) (1/3000; BioRad) at RT. After two additional washes, the membranes were developed using a chemiluminescence substrate (Clarity Western ECL Substrate, Bio-Rad) and detected for 10 s to 5 min on CL-Xposure film (Thermo Fisher Scientific) or using a Chemidoc (Bio-Rad). Films were scanned and the protein bands were quantified by measuring density via ImageJ software (National Institutes of Health, ImageJ 1.53e). GAPDH was used as the reference protein for the normalization of the proteins of interest.

### RNA extraction

Kidney organoids were manually crushed and lysed in TRIzol (Invitrogen) and mixed with chloroform (200 µl/ml Trizol). After homogenization, the samples were incubated for 15 min at RT followed by a centrifugation at 18000 RCF for 15 min at 4°C. The clear supernatant was collected, and isopropanol (500 µl/ml Trizol) was added to precipitate the RNA. The samples were homogenized by vortexing and incubated for 15–20 min at RT and centrifuged again at 4°C and 18000 RCF for 15 min. The RNA pellets were then washed several times with ethanol. After centrifugation at 4°C and 6800 RCF for 5 min, the ethanol was removed, and the pellet was air-dried before being eluted in RNAse-free water (Qiagen).

### Reverse transcription and qPCR

Reverse transcription was performed using 500 ng of RNA mixed with water, reverse transcription enzyme and iScript buffer (BioRad) in a final volume of 20 µl. The samples were heated for 5 min at 25°C for priming followed by 20 min at 46°C for reverse transcription and 1 min at 95°C for enzyme inactivation. The obtained cDNA was diluted five times in RNAse-free water, and 2 µl were prepared for qPCR with a reaction mix comprising 250 nM of each primer (Supplementary Table 2), 4 µl of water and 10 µl SYBR (Bio-Rad) Green Master Mix. The real-time PCR CFX96 (Bio-Rad) was programmed to perform 40 cycles of 2-steps amplification consisting of 10 s of denaturation at 95°C, and 60 s of combined annealing and extension at 60°C. Experiments were performed in technical duplicate and normalized to *GAPDH* as a housekeeping gene.

### Statistics

All data are expressed as the mean ± standard deviation (SD). Experiments were performed at least 3 independent times (N=3). Normality was checked using the Shapiro-Wilk normality test. Significant differences (p < 0.05) between groups were assessed using a Student T-test (2 groups) or a two-way ANOVA followed by a Tukey-Kramer post hoc ANOVA test (≥ 3 groups). Nonparametric data were analyzed using a Mann-Whitney or Kruskal-Wallis test.

## Acknowledgments

This work is supported by the partners of Regenerative Medicine Crossing Borders (RegMed XB), a public-private partnership that uses regenerative medicine strategies to cure common chronic diseases. This collaboration project is financed by the Dutch Ministry of Economic Affairs by means of the PPP Allowance made available by the Top Sector Life Sciences & Health to stimulate public-private partnerships. This project has received funding from the European Union’s Horizon 2020 research and innovation programme under the Marie Skłodowska-Curie grant agreement No 101065328. We acknowledge Ton Rabelink, Cathelijne van den Berg and Loes Wiersma from Leiden UMC for training and advice on kidney organoid cultures. We thank Nadia Roumans from the MERLN Institute for technical support regarding organoid cultures.

## Authors contribution

VJ: design of the work, data acquisition, data analysis, writing and review of manuscript AS: data acquisition, data analysis, review of the manuscript

PM: data acquisition, data analysis, review of the manuscript MEL: data acquisition, data analysis, review of the manuscript

MvG: contribution in design of the work, review of the manuscript.

VLP: conception and design of the project, data analysis, writing and review of the manuscript, fund raising

## Declaration

A patent application related to this work was submitted.

---

**Figure S1:**
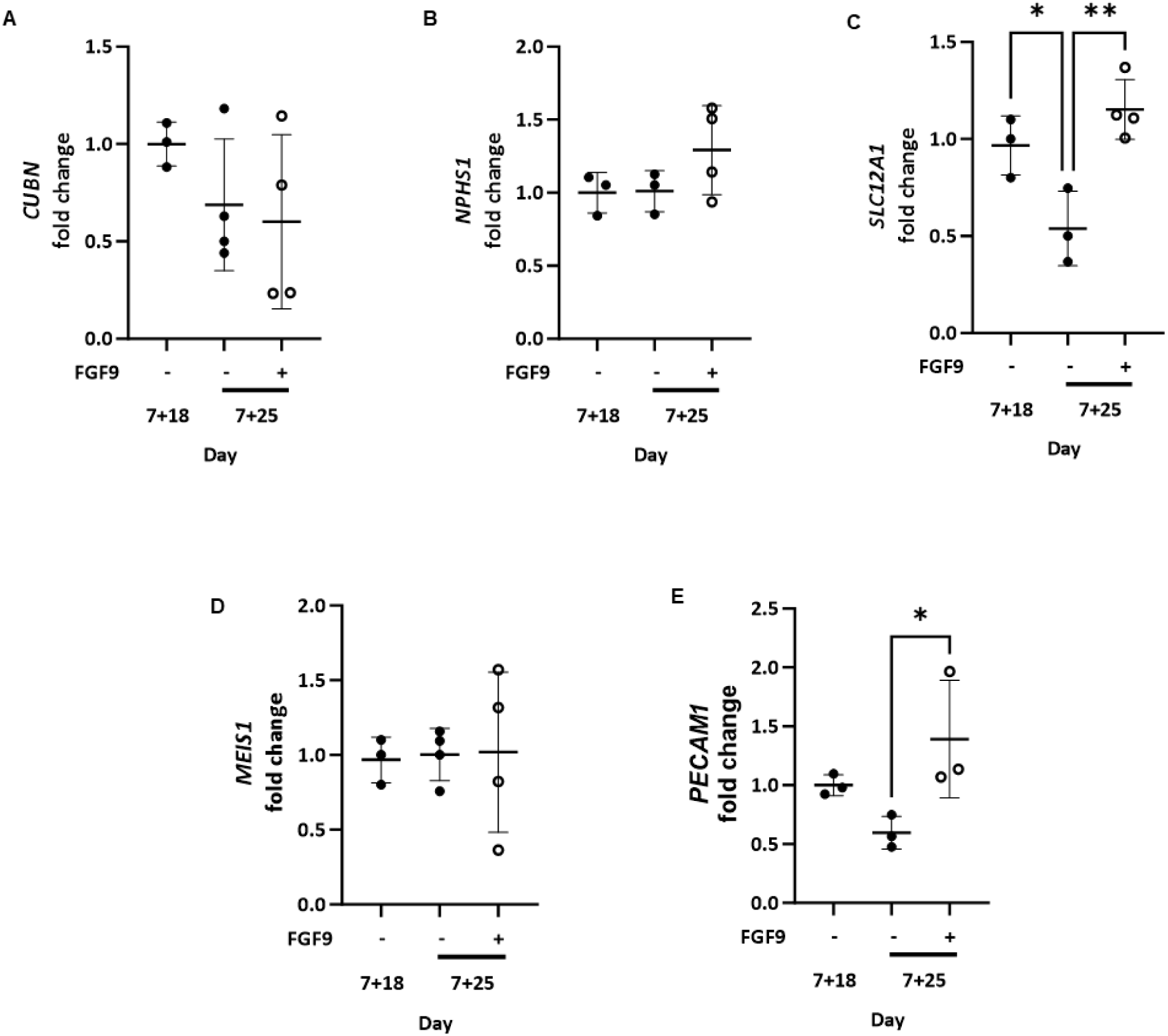
FGF9 treatment does not negatively affect renal structures. (A-D) Gene expression of markers of renal structures, *CUBN, NPHS1, SLC12A1* and stromal population *MEIS1* were assessed by qPCR and shown as -fold change compared to expression at day 7+18. *\*p* < 0.05; *\*\*p* < 0.01; from 3-4 samples. (E) FGF9 treatment (+) upregulates *PECAM1* expression compared to control (−) organoids at day 7+25. *PECAM1* expression assessed using qPCR shown as -fold change compared to expression in untreated organoids at day 7+ 18. *\*p* < 0.05 from 3 samples.

**Figure S2:**
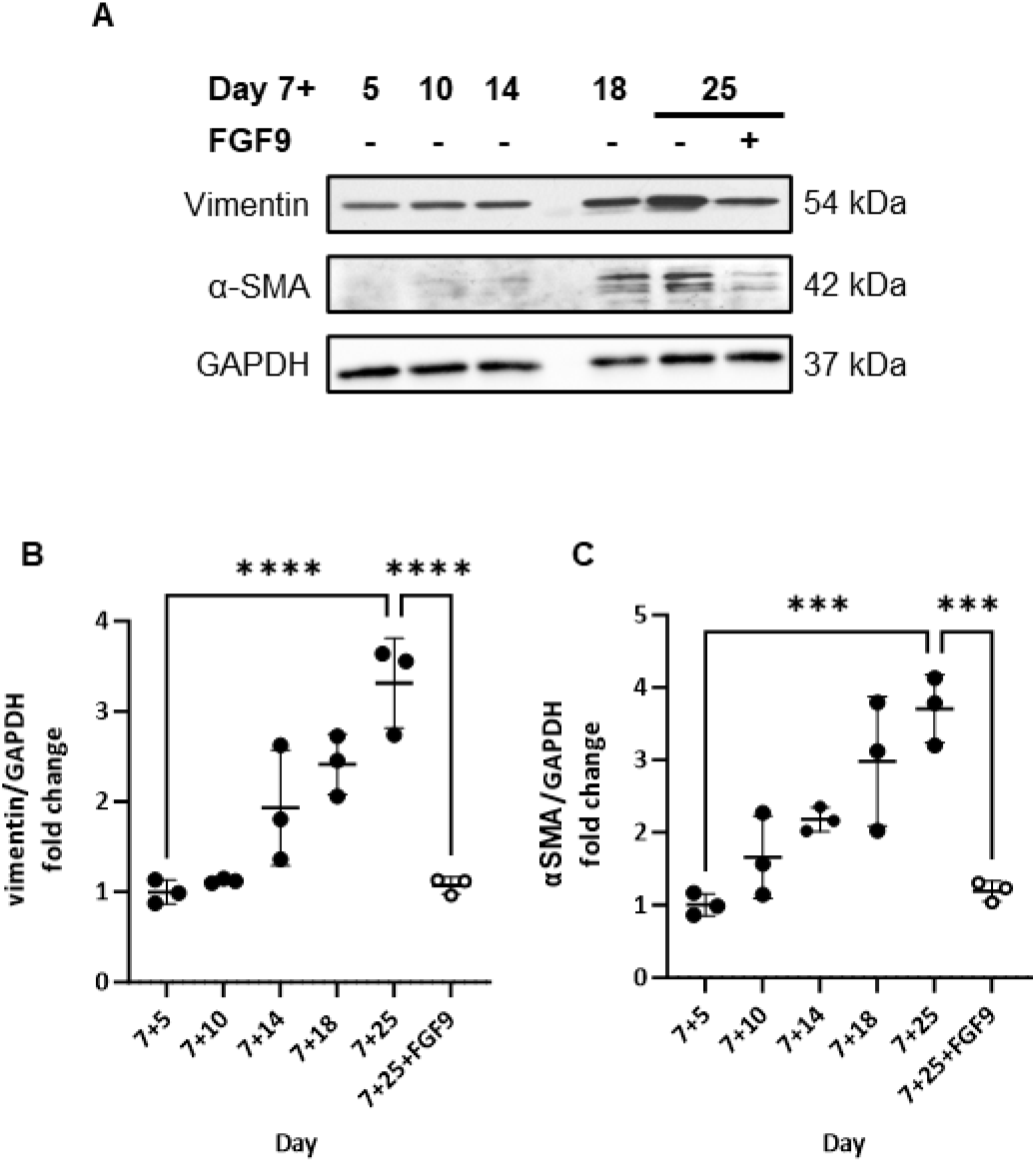
FGF9-treated organoids express lower levels of EMT markers at day 7+25. (A) Western immunoblotting showed lower levels of EMT markers vimentin and α-SMA in FGF9-treated (+) organoids compared to control (−) organoids at day 7+25. Notably, vimentin and α-SMA progressively increased from day 7+5, 7+10, 7+14, 7+18 and 7+25 in control organoids. GADPH levels are shown as loading controls. (B-C) Quantification of vimentin (B) and α-SMA (C) levels at day 7+5, 7+10, 7+14, 7+18 and 7+25 showed significant increases in untreated organoids, which were ameliorated by FGF9 treatment (7+25+FGF9). Protein levels are expressed as - fold change relative to day 7+5. *** *p* < 0.001; **** *p* < 0.0001 from 3 samples each.

**Figure S3.**
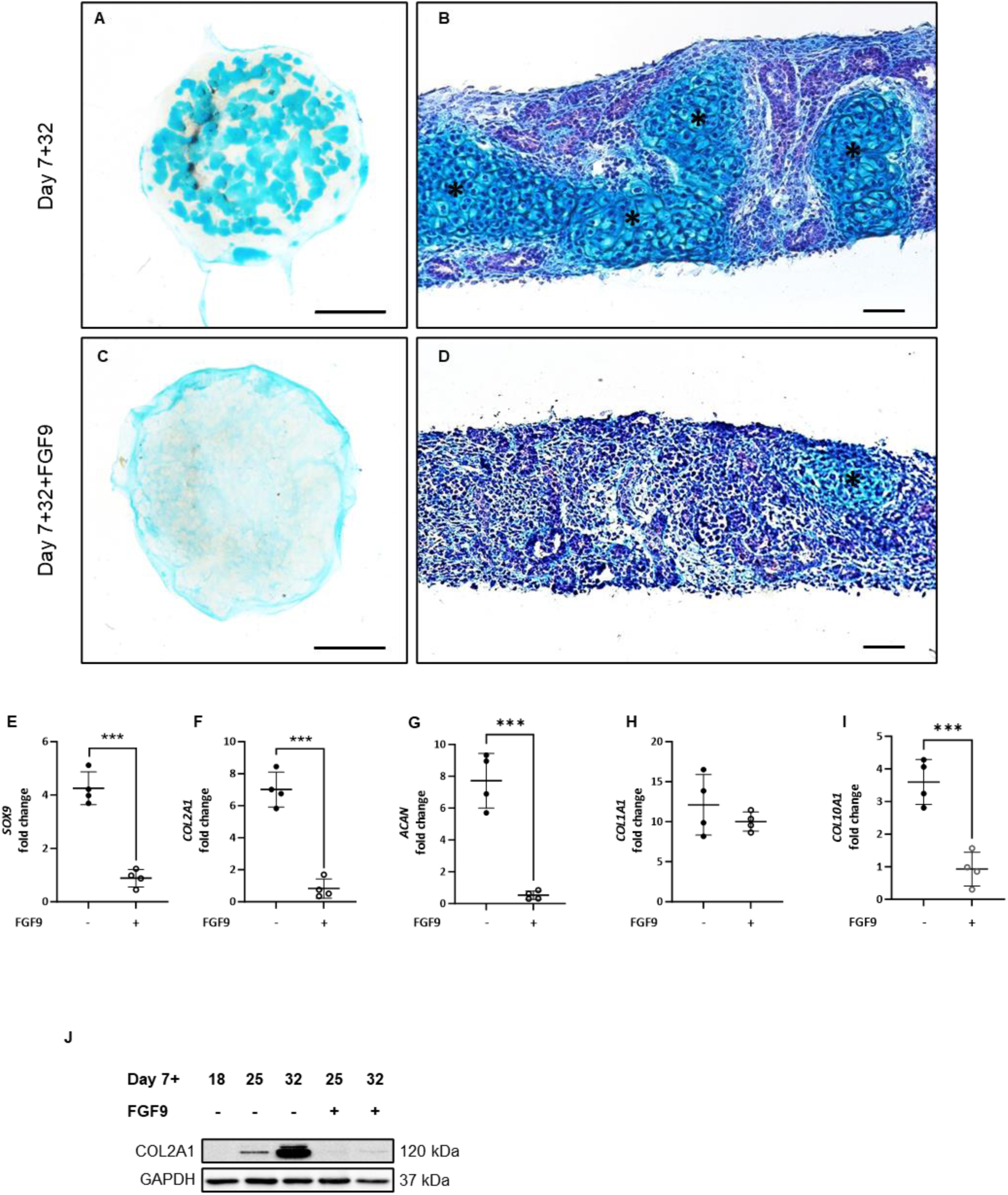
FGF-9 treatment delays the appearance of cartilage in iPSC-derived kidney organoids. (A-D) Cartilage (asterisks) stained with Alcian blue in whole organoids (left images, scale bars represent 1000 μm) and on cryosectionos (rights images, scale bars represent 50 μm) was less abundant with FGF9 treatment but the appearances of small islands of cartilage were visible (bottom row). (E-I) FGF9 treatment (+) significantly decreased four of five markers of chondrongenesis in kidney organoids at day 7+32 compared to control (−) organoids. Gene expression assessed by qPCR and show as -fold change compared to expresssin in untreated organoids at day 7+18. ***p < 0.001 from 4 samples. (J) Western blotting COL2A1 protein in control (−) kidney organoids showed increased expression over time which was abrogated with FGG9 treatment (+). GADPH levels shown as loading controls.

**Supplementary Table 1:** Primary and secondary antibodies used for immunofluorescence.

| **Primary antibodies** | **Species** | **Company** | **Reference** | **Dilution** |
| --- | --- | --- | --- | --- |
| Nephrin 1 | Sheep | R&D Systems | AF4269 | 1/300 |
| E-cadherin | Mouse | Beckton Dickinson | BD 610182 | 1/300 |
| SLC12A1 | Rabbit | Sigma-Aldrich | HPA014967 | 1/300 |
| MEIS1/2/3 | Mouse | Santa Cruz Biotechnology | sc-101850 | 1/300 |
| CD31 | Mouse | Abcam | ab24590 | 1/300 |
| Lotus lectin biotin conjugated (LTL) | / | Brunschwig | B-1325 | 1/1000 |
| Aquaporin 2 | Rabbit | Abcam | ab199975 | 1/300 |
| **Secondary antibodies** |  | **Company** | **Reference** | **Dilution** |
| Alexa Fluor donkey anti-sheep 488 |  | Thermo Fisher Scientific | A-11015 | 1/400 |
| Alexa Fluor goat anti-mouse 568 |  | Thermo Fisher Scientific | A-11031 | 1/400 |
| Alexa Fluor goat anti-rabbit 568 |  | Thermo Fisher Scientific | A-10042 | 1/400 |
| Alexa Fluor goat anti-mouse 488 |  | Thermo Fisher Scientific | A-11029 | 1/400 |
| Alexa Fluor goat anti-sheep 647 |  | Thermo Fisher Scientific | A-21448 | 1/400 |
| Streptavidin Alexa Fluor 647 conjugate |  | Thermo Fisher Scientific | S-21374 | 1/1000 |

**Supplementary Table 2:**
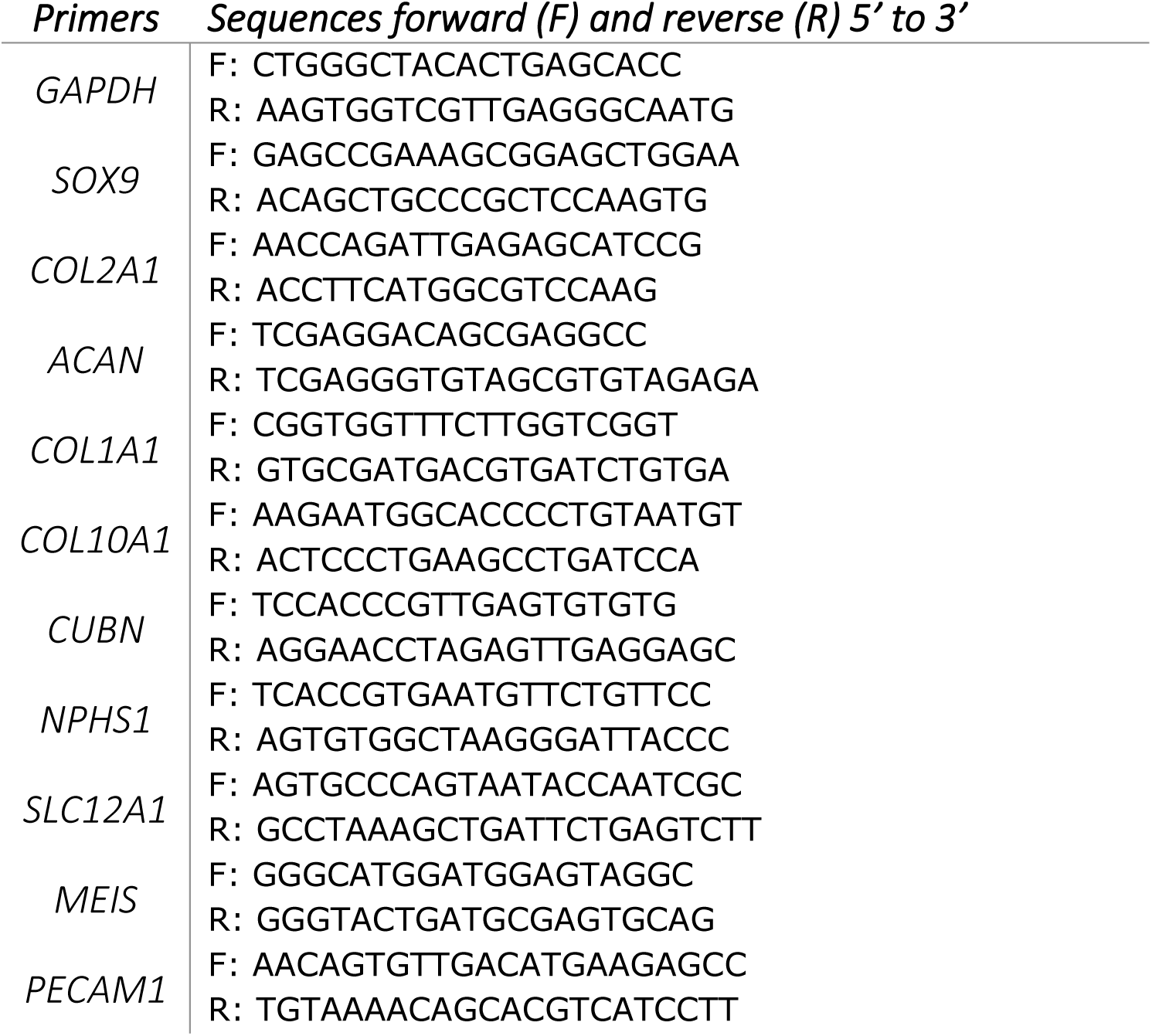
Primers used in qPCR.

## References

1 Ghane Shahrbaf, F. & Assadi, F. Drug-induced renal disorders. J Renal Inj Prev 4, 57–60, doi:10.12861/jrip.2015.12 (2015).

2 Benam, K. H. et al. Engineered in vitro disease models. Annu Rev Pathol 10, 195–262, doi:10.1146/annurev-pathol-012414-040418 (2015).

3 Fatehullah, A., Tan, S. H. & Barker, N. Organoids as an in vitro model of human development and disease. Nat Cell Biol 18, 246–254, doi:10.1038/ncb3312 (2016).

4 Takasato, M. et al. Kidney organoids from human iPS cells contain multiple lineages and model human nephrogenesis. Nature 526, 564–568, doi:10.1038/nature15695 (2015).

5 Geuens, T. et al. Thiol-ene cross-linked alginate hydrogel encapsulation modulates the extracellular matrix of kidney organoids by reducing abnormal type 1a1 collagen deposition. Biomaterials 275, 120976, doi:10.1016/j.biomaterials.2021.120976 (2021).

6 van den Berg, C. W., Koudijs, A., Ritsma, L. & Rabelink, T. J. In Vivo Assessment of Size-Selective Glomerular Sieving in Transplanted Human Induced Pluripotent Stem Cell-Derived Kidney Organoids. J Am Soc Nephrol 31, 921–929, doi:10.1681/ASN.2019060573 (2020).

7 Forbes, T. A. et al. Patient-iPSC-Derived Kidney Organoids Show Functional Validation of a Ciliopathic Renal Phenotype and Reveal Underlying Pathogenetic Mechanisms. Am J Hum Genet 102, 816–831, doi:10.1016/j.ajhg.2018.03.014 (2018).

8 Cruz, N. M. et al. Organoid cystogenesis reveals a critical role of microenvironment in human polycystic kidney disease. Nat Mater 16, 1112–1119, doi:10.1038/nmat4994 (2017).

9 Romero-Guevara, R., Ioannides, A. & Xinaris, C. Kidney Organoids as Disease Models: Strengths, Weaknesses and Perspectives. Front Physiol 11, 563981, doi:10.3389/fphys.2020.563981 (2020).

10 Low, J. H. et al. Generation of Human PSC-Derived Kidney Organoids with Patterned Nephron Segments and a De Novo Vascular Network. Cell Stem Cell 25, 373–387 e379, doi:10.1016/j.stem.2019.06.009 (2019).

11 Schumacher, A. et al. Enhanced Microvasculature Formation and Patterning in iPSC-Derived Kidney Organoids Cultured in Physiological Hypoxia. Front Bioeng Biotechnol 10, 860138, doi:10.3389/fbioe.2022.860138 (2022).

12 Kim, J. W. et al. Kidney Decellularized Extracellular Matrix Enhanced the Vascularization and Maturation of Human Kidney Organoids. Adv Sci (Weinh) 9, e2103526, doi:10.1002/advs.202103526 (2022).

13 Geuens, T., van Blitterswijk, C. A. & LaPointe, V. L. S. Overcoming kidney organoid challenges for regenerative medicine. NPJ Regen Med 5, 8, doi:10.1038/s41536-020-0093-4 (2020).

14 Wu, H. et al. Comparative Analysis and Refinement of Human PSC-Derived Kidney Organoid Differentiation with Single-Cell Transcriptomics. Cell Stem Cell 23, 869–881 e868, doi:10.1016/j.stem.2018.10.010 (2018).

15 Nam, S. A. et al. Graft immaturity and safety concerns in transplanted human kidney organoids. Exp Mol Med 51, 1–13, doi:10.1038/s12276-019-0336-x (2019).

16 Bantounas, I. et al. Generation of Functioning Nephrons by Implanting Human Pluripotent Stem Cell-Derived Kidney Progenitors. Stem Cell Reports 10, 766–779, doi:10.1016/j.stemcr.2018.01.008 (2018).

17 Li, J. & Dong, S. The Signaling Pathways Involved in Chondrocyte Differentiation and Hypertrophic Differentiation. Stem Cells Int 2016, 2470351, doi:10.1155/2016/2470351 (2016).

18 Tang, J., Liu, N. & Zhuang, S. Role of epidermal growth factor receptor in acute and chronic kidney injury. Kidney Int 83, 804–810, doi:10.1038/ki.2012.435 (2013).

19 Kumar, S. et al. Sox9 Activation Highlights a Cellular Pathway of Renal Repair in the Acutely Injured Mammalian Kidney. Cell Rep 12, 1325–1338, doi:10.1016/j.celrep.2015.07.034 (2015).

20 Bobick, B. E., Tuan, R. S. & Chen, F. H. The intermediate filament vimentin regulates chondrogenesis of adult human bone marrow-derived multipotent progenitor cells. J Cell Biochem 109, 265–276, doi:10.1002/jcb.22419 (2010).

21 Lefebvre, V. & Dvir-Ginzberg, M. SOX9 and the many facets of its regulation in the chondrocyte lineage. Connect Tissue Res 58, 2–14, doi:10.1080/03008207.2016.1183667 (2017).

22 Barak, H. et al. FGF9 and FGF20 maintain the stemness of nephron progenitors in mice and man. Dev Cell 22, 1191–1207, doi:10.1016/j.devcel.2012.04.018 (2012).

23 Walker, K. A., Sims-Lucas, S. & Bates, C. M. Fibroblast growth factor receptor signaling in kidney and lower urinary tract development. Pediatr Nephrol 31, 885–895, doi:10.1007/s00467-015-3151-1 (2016).

24 Correa, D. et al. Sequential exposure to fibroblast growth factors (FGF) 2, 9 and 18 enhances hMSC chondrogenic differentiation. Osteoarthritis Cartilage 23, 443–453, doi:10.1016/j.joca.2014.11.013 (2015).

25 Schumacher, A., Joris, V., Griensven, M. v. & LaPointe, V. Ultrastructural comparison of human kidney organoids and human fetal kidneys reveals features of hyperglycemic culture. bioRxiv, 2023.2003.2027.534124, doi:10.1101/2023.03.27.534124 (2023).

26 Schumacher, A. et al. Defining the variety of cell types in developing and adult human kidneys by single-cell RNA sequencing. NPJ Regen Med 6, 45, doi:10.1038/s41536-021-00156-w (2021).

27 Huh, S. H., Ha, L. & Jang, H. S. Nephron Progenitor Maintenance Is Controlled through Fibroblast Growth Factors and Sprouty1 Interaction. J Am Soc Nephrol 31, 2559–2572, doi:10.1681/ASN.2020040401 (2020).

28 Zhang, X., Weng, M. & Chen, Z. Fibroblast Growth Factor 9 (FGF9) negatively regulates the early stage of chondrogenic differentiation. PLoS One 16, e0241281, doi:10.1371/journal.pone.0241281 (2021).

29 Li, L., Zhang, S., Li, H. & Chou, H. FGFR3 promotes the growth and malignancy of melanoma by influencing EMT and the phosphorylation of ERK, AKT, and EGFR. BMC Cancer 19, 963, doi:10.1186/s12885-019-6161-8 (2019).

30 Wang, H. & Unternaehrer, J. J. Epithelial-mesenchymal Transition and Cancer Stem Cells: At the Crossroads of Differentiation and Dedifferentiation. Dev Dyn 248, 10–20, doi:10.1002/dvdy.24678 (2019).

31 Ruiter, F. A. A. et al. Soft, Dynamic Hydrogel Confinement Improves Kidney Organoid Lumen Morphology and Reduces Epithelial-Mesenchymal Transition in Culture. Adv Sci (Weinh) 9, e2200543, doi:10.1002/advs.202200543 (2022).

32 Behr, B., Leucht, P., Longaker, M. T. & Quarto, N. Fgf-9 is required for angiogenesis and osteogenesis in long bone repair. Proc Natl Acad Sci U S A 107, 11853–11858, doi:10.1073/pnas.1003317107 (2010).

33 Frontini, M. J. et al. Fibroblast growth factor 9 delivery during angiogenesis produces durable, vasoresponsive microvessels wrapped by smooth muscle cells. Nat Biotechnol 29, 421–427, doi:10.1038/nbt.1845 (2011).

34 Kwon, T. H., Frokiaer, J. & Nielsen, S. Regulation of aquaporin-2 in the kidney: A molecular mechanism of body-water homeostasis. Kidney Res Clin Pract 32, 96–102, doi:10.1016/j.krcp.2013.07.005 (2013).

35 Kwon, T. H. et al. Altered expression of renal AQPs and Na(+) transporters in rats with lithium-induced NDI. Am J Physiol Renal Physiol 279, F552–564, doi:10.1152/ajprenal.2000.279.3.F552 (2000).

36 Fan, Y. et al. Overexpression of aquaporin 2 in renal tubular epithelial cells alleviates pyroptosis. Transl Androl Urol 10, 2340–2350, doi:10.21037/tau-21-71 (2021).

37 Lai, M. S., Cheng, Y. S., Chen, P. R., Tsai, S. J. & Huang, B. M. Fibroblast growth factor 9 activates akt and MAPK pathways to stimulate steroidogenesis in mouse leydig cells. PLoS One 9, e90243, doi:10.1371/journal.pone.0090243 (2014).

